# Comparative Landscape of Small RNAs in Tissue and Liquid Biopsies for Liver Transplant Outcomes

**DOI:** 10.1101/2025.09.16.676609

**Authors:** Monima Anam, Christine Watkins, Grace Rucker, Katherine Marlow, Matthew Khalil, Murat Dogan, James Eason, Jason Vanatta, Corey Eymard, Cem Kuscu, Zhangli Su, Canan Kuscu

## Abstract

**Background:** Ischemia-reperfusion injury (IRI) is an inevitable consequence of liver transplantation, arising during donor organ procurement and reoxygenation. Severe IRI is a leading contributor to early allograft dysfunction (EAD), a post-transplant complication associated with reduced graft survival. Current postoperative biomarkers provide limited time for intervention, highlighting a need to identify preoperative biomarkers of IRI. Meanwhile, tRNA fragments (tRFs) have emerged as novel biomarkers in various diseases but remain unexplored in the context of liver transplant.

**Results:** We performed small RNA sequencing on 96 paired donor liver biopsies from 48 patients to investigate IRI-associated transcript changes. In parallel, 161 donor liver perfusates were analyzed as a non-invasive surrogate for tissue. Across samples, microRNAs (miRNAs) and tRFs were the most abundant. Perfusate expression strongly correlated with biopsies, supporting their value as a non-invasive source of small RNAs. Comparison between post-reperfusion and pre-implantation biopsies revealed that IRI reprogrammed tRF expression. Stratification by clinical outcome showed that patients who developed EAD exhibited specific small RNA signatures in both biopsy and perfusate. Receiver operating characteristic (ROC) analysis revealed a miRNA-based model that achieved an AUC of 0.772, outperforming donor risk index alone (AUC = 0.665), representing a 10.7% increase in discriminative capacity.

**Conclusions:** These results are the first to establish tRFs as IRI-responsive biomolecules abundant in both donor liver tissue and non-invasive perfusate. In particular, various small RNAs emerged as promising candidate biomarkers for early detection of EAD. These results lay the foundation to further investigate the prognostic utility of tRFs/miRNAs in liver transplantation.

## Background

Ischemia-reperfusion injury (IRI) is an inevitable consequence of solid organ transplantation, initiated during donor organ procurement and cold storage, and exacerbated upon reperfusion in the recipient. The sudden reoxygenation of ischemic tissue induces a complex interplay of oxidative stress, metabolic disturbances, sterile inflammation, and cell death pathways that impair graft integrity (1). Severe IRI is a principal contributor to early allograft dysfunction (EAD), a clinical syndrome presenting within the first week post liver transplant, associated with increased morbidity, primary nonfunction, and diminished long-term graft survival (2). Current diagnostic criteria for EAD primarily rely on postoperative serum biochemical markers such as alanine aminotransferase (ALT) and aspartate aminotransferase (AST), which provide limited mechanistic insight and insufficient lead time for intervention (3). This underscores the need for high-resolution molecular biomarkers capable of detecting early transcriptional and post-transcriptional responses to IRI in graft tissue and accessible biofluids, thereby enabling predictive and mechanism-informed monitoring after liver transplantation.

Small RNAs are central regulators of post-transcriptional gene expression, modulating mRNA stability, translation, and chromatin organization across diverse biological contexts. Beyond well-characterized microRNAs (miRNAs) and piRNAs, transfer RNA-derived fragments (tRFs) and stress-induced tRNA halves (tiRs) have recently emerged as abundant and functionally versatile small RNA species generated by precise cleavage of precursor or mature tRNAs (4). Their biogenesis is strongly induced by cellular stressors, including oxidative injury, nutrient deprivation, hypoxia, and immune activation (5,6). Functionally, tRFs repress global protein synthesis, promote stress granule assembly, displace messenger RNAs from RNA-binding proteins such as YBX1, and participate in RNA silencing pathways via Argonaute association (7–10). A recent publication also demonstrated that hypoxia induced tRF promotes cellular homeostasis and protects against kidney injury by enhancing autophagic flux (11). Through these mechanisms, tRFs act as dynamic modulators of cellular adaptation to stress, influencing survival, metabolism, and signaling, positioning them as promising molecular sentinels of stress phenotypes in both experimental and clinical settings (12,13).

Recent advances in transcriptomic profiling have identified small RNA signatures as a promising class of molecular biomarkers for disease diagnosis, prognosis, and therapeutic monitoring. Their inherent stability in biofluids, resistance to RNase degradation, and ability to reflect dynamic physiological states make them well-suited for minimally invasive diagnostics (14,15). miRNAs have been explored as potential biomarkers in kidney and liver transplant (16–18). In liver transplantation, liver-specific miR-122 detected in recipient serum was shown to be correlated with ALT and AST levels, thus IRI severity and EAD (19). Furthermore, donor organ preservation solution (perfusate) has emerged as an attractive biospecimen for biomarker discovery, capturing molecules released directly from the donor liver during storage and perfusion (20,21). While these investigations provide important proof-of-concepts, they focused on targeted RNA species rather than unbiased transcriptome-wide profiling. Consequently, the broader RNA repertoire within perfusate, including stress-responsive small RNAs such as tRFs and tiRs, remains largely unexplored. Here, we report the first transcriptome-wide small RNA profiling of 96 paired liver transplant biopsies from 48 patients alongside perfusate samples from 161 donor organs, aimed at uncovering mechanistically linked early non-invasive biomarkers of graft function that complement tissue-based assays.

## Results

### Small RNAs and tRFs are abundantly expressed in donor liver transplant biopsy samples

To characterize small RNAs present in donor liver tissue and determine whether they change from pre– to post-transplantation, we collected paired donor liver biopsy samples (n = 48) at two timepoints: pre-implantation (L1) and post-reperfusion (L2), followed by small RNA sequencing analysis (**Fig. 1A**). Principal component analysis (PCA) of all 96 samples revealed distinct separation between L1 and L2 small RNA expression profiles, with PC1 and PC2 accounting for 19% and 12% of the variance, respectively (**Fig. 1B).** A variety of small RNA species were detected through sequencing; however, miRNAs comprised the largest proportion of mapped reads, followed by tRFs (**Fig. 1C**). Four tRF subtypes dominated the tRF landscape: tiR5, tRF5, tRF3CCA, and tRF1 species (**Fig. 1D and E**). tiR5s are known as 5’ tRNA halves, which are typically 30–35 nucleotides (nt) in length, while tRF5s are shorter 5’ tRNA fragments that range from 15–29 nt long. tRF3CCAs are 3’ tRNA fragments that retain 3’ CCA tails, a feature of mature tRNAs. While tRF1s are the remnant 3’ trailer sequences that are cleaved from precursor tRNAs to form mature tRNAs. These four tRF subtypes represented the majority of mapped tRF reads across samples. Additionally, tRFs derived from Glu– and Gly-carrying parental tRNAs were highly enriched relative to other amino acid isotypes (**Fig. S1A**). Furthermore, multiple specific tRF species rivaled the expression of the topmost expressed miRNAs, such as Glu CTC tiR5 and Gly GCC tiR5, which were the fourth– and sixth-most highly expressed small RNAs across samples (**Fig. S1A-B**). Overall, we observe that tRFs are steadily abundant across L1 and L2 samples, serving as promising candidates for further investigation.

**Figure 1.**
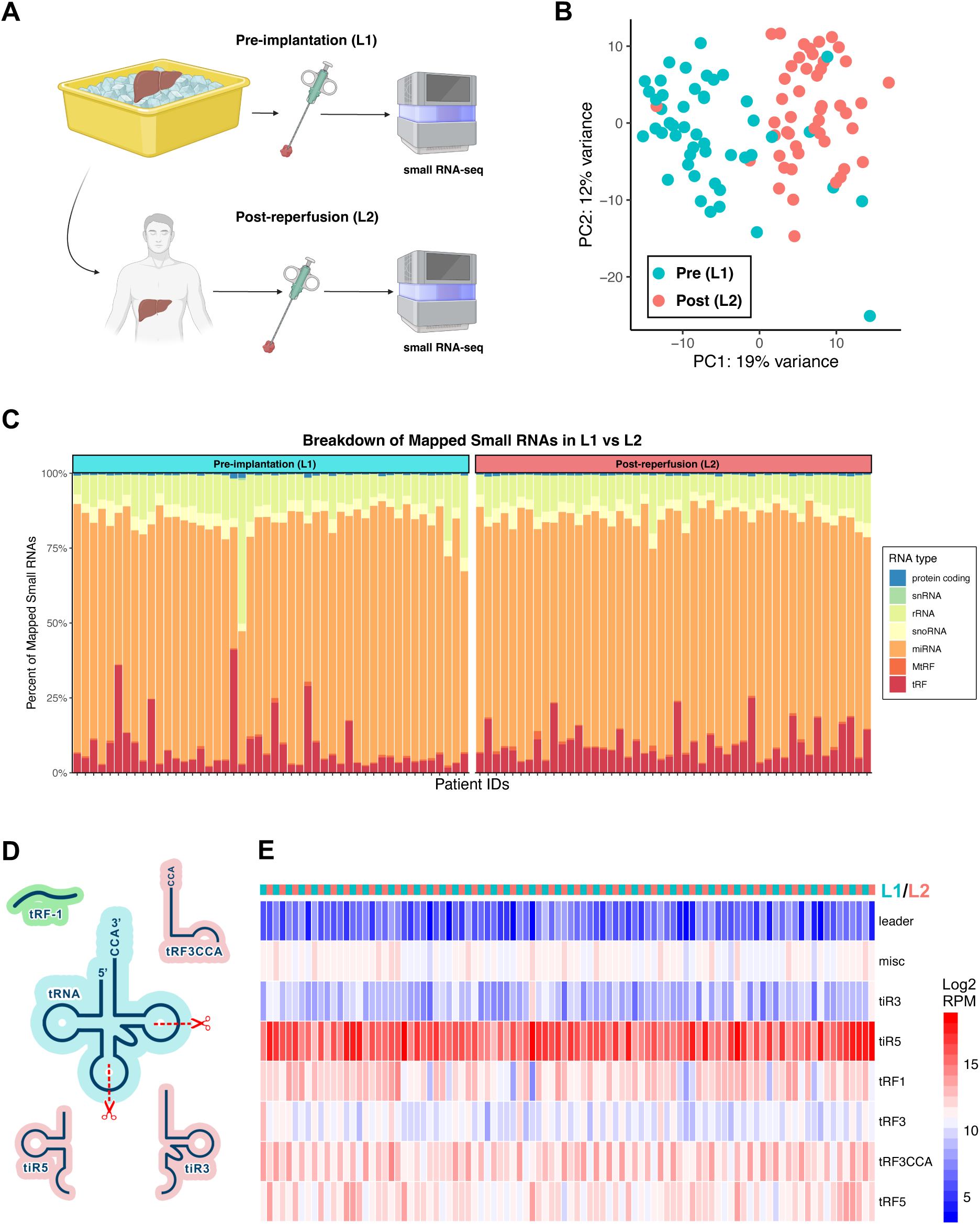
Abundance of tRNA fragments (tRFs) in 96 donor liver transplant biopsy samples. **(A)** Paired liver biopsy samples were obtained from donor livers at two time points: pre-implantation (L1, n = 48) and post-reperfusion (L2, n = 48), followed by small RNA sequencing. **(B)** Principal component analysis (PCA) of small RNA sequencing data reveals distinct clustering between L1 and L2 small RNA profiles. **(C)** Stacked bar plot displaying the proportion of mapped reads for each small RNA type in L1 and L2 samples. For each patient (x-axis), the bars represent the relative proportion of total reads (y-axis) assigned to each small RNA type. Across samples, miRNAs comprised the largest majority of mapped reads, followed by tRFs. **(D)** Schematic illustrating four highlighted tRF subtypes. The tRF nomenclature echoes the region of tRNA from which tRFs are derived. The fragments shown are examples of each tRF subtype and do not represent all possible variants. tRF-1 species (green) are derived from precursor tRNAs (not pictured), while tiR5, tiR3, and tRF3CCA species (pink) are cleaved from mature tRNAs (blue). **(E)** Heatmap displaying the relative expression (Log2 RPM) of different tRF subtypes across L1 and L2 samples. Overall, biopsy samples showed abundant tRF expression.

### Comparison of L2 vs. L1 expression profiles reveals IRI-associated tRF changes

We next examined small RNAs differentially expressed in response to IRI by comparing L2 and L1 samples, as L2 samples were collected post-reperfusion and therefore serve as a proxy for transplant-associated IRI. DESeq2 analysis of paired L2 vs. L1 small RNA profiles revealed 182 differentially expressed small RNAs, including 116 tRFs and 66 miRNAs (**Supplementary Table 1**). Of these, 102 (58 tRFs, 44 miRNAs) were upregulated and 80 (58 tRFs, 22 miRNAs) were downregulated in L2 compared to L1 (**Fig. 2A and B**). Strikingly, tRFs of the same subtype displayed uniform directional changes. Among the most abundant tRF classes, differentially expressed tiR5s were consistently enriched in L2, while significant tRF3CCA species were consistently depleted in L2. Similar trends were observed for differentially expressed tRF1 and tRF5 species, with the majority showing upregulation following IRI (**Fig. 2B**). Boxplots of select tiR5 and tRF3CCA species likewise exemplify and highlight this trend (**Fig. 2C-E**). Together, these findings indicate that IRI is associated with subtype-specific changes in tRF expression, suggesting that tRF dysregulation may be a characteristic feature of IRI.

**Figure 2.**
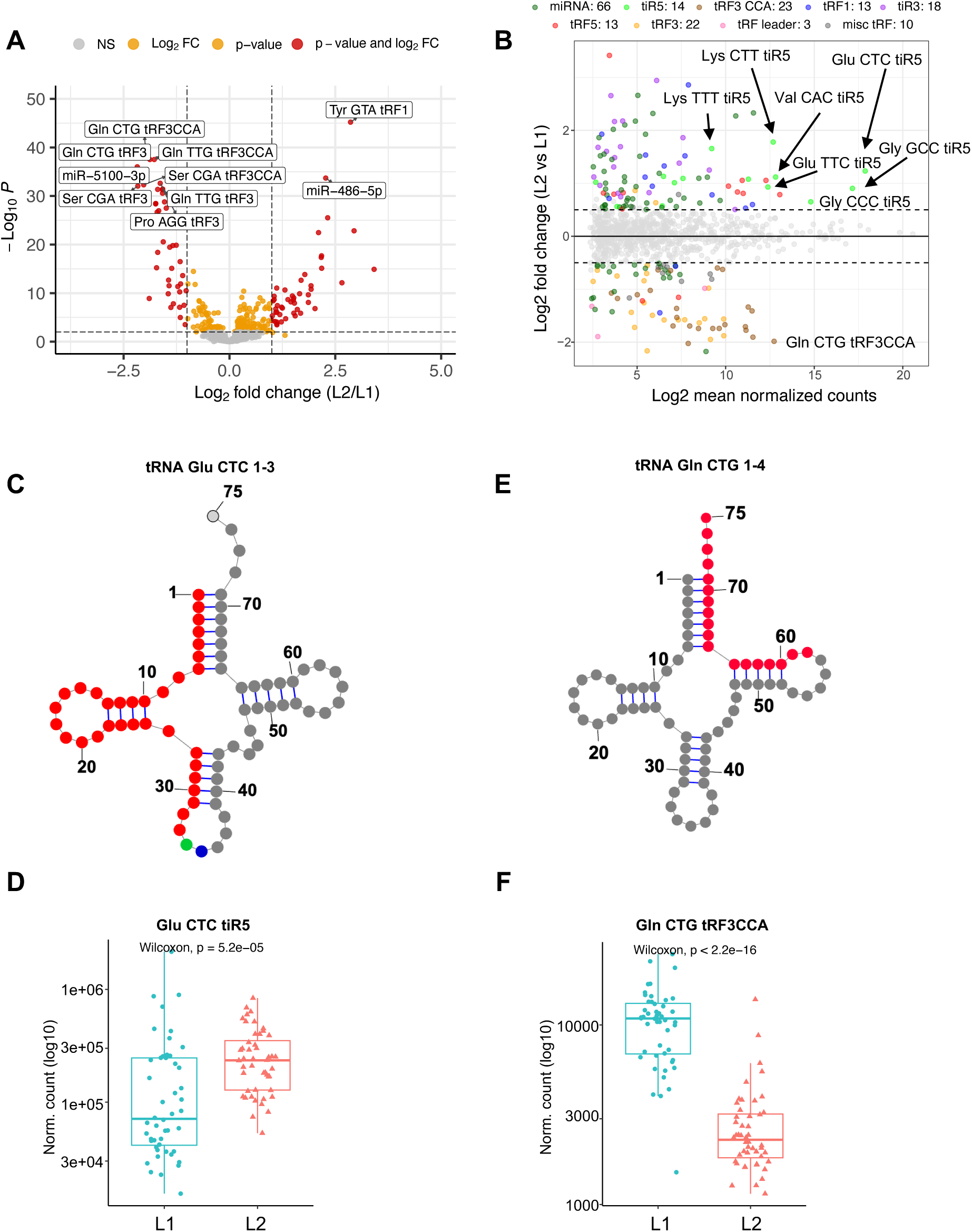
Significant tRF alterations associated with ischemia-reperfusion injury (IRI) during liver transplantation. **(A)** Volcano plot of differentially expressed small RNAs between paired L2 and L1 samples, identified using DESeq2. The x-axis shows the log2 fold change (L2 vs. L1), and the y-axis shows statistical significance (−log10 adjusted p-value). The top 10 most significantly altered small RNAs are labeled, many being tRF3CCA species downregulated in L2. **(B)** MA plot of differentially expressed small RNAs (L2 vs. L1) identified using DESeq2 (p. adj < 0.05, |Log2FoldChange| > 0.5). The x-axis represents gene expression (log2 normalized counts), and the y-axis shows the log2 fold change. Differentially expressed RNAs are color-coded by RNA type (tRF subtype or miRNA). Select highly abundant, significantly altered tRFs are labeled, including many tiR5 species that are upregulated in L2. **(C)** Primary and secondary structure of parental tRNA-Glu-CTC-1-3, with its most abundant tiR5 fragment highlighted in red (positions 1-35). Cleavage sites that generate the second and third most abundant Glu CTC tiR5 isoforms from tRNA-Glu-CTC are marked in green and blue. **D)** Boxplot of Glu CTC tiR5 expression in L1 (n = 48) vs. L2 (n = 48) samples, showing significant upregulation in L2, consistent with IRI-associated induction. **(E)** Primary and secondary structure of parental tRNA-Gln-CTG-1-4, with its corresponding tRF3CCA fragment highlighted in red (positions 58–75). **(F)** Boxplot of Gln CTG tRF3CCA expression in L1 (n = 48) vs. L2 (n = 48) samples, showing significant downregulation in L2, indicative of IRI-associated suppression.

### IRI-associated tRF changes are stratified by transplant outcome

Next, we stratified patients by early transplant outcome to assess whether IRI-associated tRF changes were uniquely associated with graft dysfunction. Transplant outcomes were based on day 1 AST values (2), where high AST values (>2000) corresponded to early allograft dysfunction (EAD, n = 18) and moderate/low AST values corresponded to normal function (NF, n = 30) **(Fig. 3A and S2A)**. DESeq2 analysis of L2 vs. L1 in each outcome group (EAD or NF) revealed group-dependent regulation of tRFs. In the EAD group, L2 biopsies showed unique upregulation of numerous tiR5 species not observed in NF samples **(Fig. 3B and C)**. In contrast, the NF group exhibited unique upregulation of multiple tRF1 species in L2 vs. L1 comparisons **(Fig. 3B, S2C and D)**. Notably, no tiR5 species were downregulated in L2 in either group **(Fig. S2E)**. Downregulated RNAs in L2 were largely shared between groups and consisted primarily of tRF3CCA species **(Fig. S2E and F)**. The complete comparison is also included in **Supplementary Table 1.** Overall, we observe that post-IRI tRF changes are uniquely dysregulated in EAD and NF grafts, suggesting that tRFs may be prognostically linked to graft outcome.

**Figure 3.**
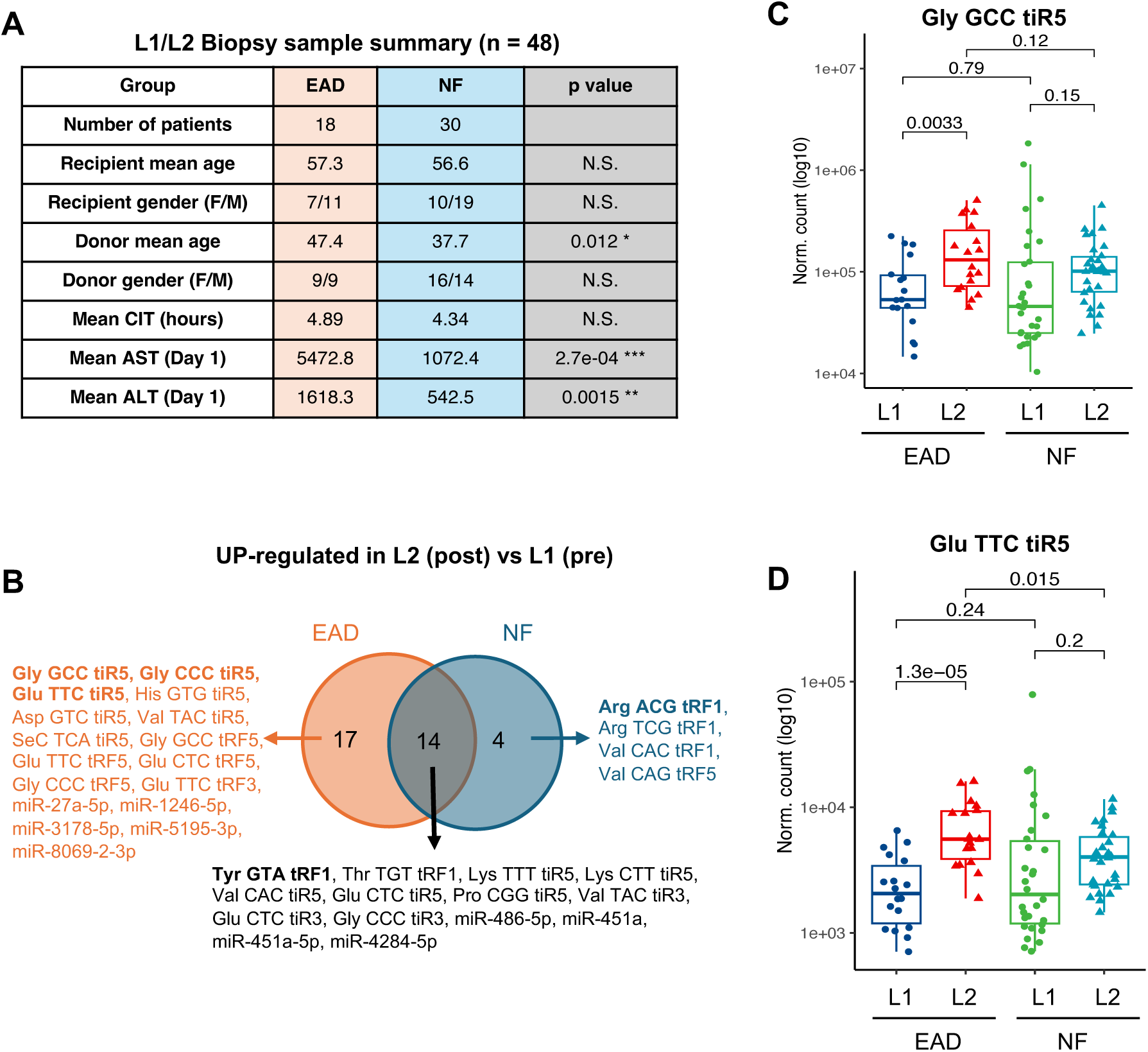
IRI-associated tRF alterations are transplant outcome specific. **(A)** Clinical and demographic characteristics of the patient cohort, stratified by transplant outcome (EAD or NF), which was classified based on day 1 post-transplant aspartate aminotransferase (AST) levels. Cold ischemia time (CIT), the duration the donor liver remained on ice prior to transplantation, averaged 4.6 hours across groups. **(B)** Differential expression analysis of paired L2 vs. L1 samples was performed separately within EAD and NF groups (p. adj < 0.05). The Venn diagram displays small RNAs upregulated in L2 relative to L1 for each group. In the EAD group (n = 18), uniquely upregulated RNAs were predominantly tiR5 species, whereas in the NF group (n = 30), uniquely upregulated RNAs were mainly tRF1 species. A subset of tiR5 and tRF1 species was commonly upregulated in both groups, alongside other tRF and miRNA species. **(C)** and **(D)** Boxplots showing normalized expression (log10 transformed) of representative tiR5 species (Gly-GCC and Glu-TTC), stratified by transplant outcome (EAD vs. NF) and biopsy timepoint (L2 vs. L1). Statistical comparisons were performed using Wilcoxon rank-sum tests; p-values are indicated above comparisons. Both panels exemplify the EAD-specific upregulation of tiR5s, showing significant upregulation in L2 compared to L1 exclusively in EAD samples (Student’s t-test).

### Comparison of EAD vs. NF small RNA profiles reveals transplant outcome-associated small RNA changes, independent of IRI

To evaluate whether small RNA profiles differed between EAD and NF grafts, we performed differential expression analysis (EAD vs. NF) separately within L1 and L2 samples, including batch correction. This approach allowed us to identify small RNAs associated with transplant outcome independent of IRI-induced changes. DESeq2 analysis revealed multiple differentially expressed small RNAs that were shared across EAD vs. NF comparisons (**Supplementary Table 2**). Specifically, miR-452-5p, miR-224-5p, and miR-296-3p were significantly upregulated in EAD samples compared to NF samples (**Fig. 4A–E and Fig. S3A**). In contrast, two small RNAs were consistently downregulated in EAD, including Gly TCC tRF3CCA and miR-1247-5p (**Fig. S3B**). To understand potential biological functions of EAD-associated miRNAs, we performed pathway enrichment analysis of predicted mRNA targets of the three miRNAs consistently upregulated in EAD samples. Network analysis revealed that miR-224-5p, miR-452-5p, and miR-296-3p share overlapping target gene networks (**Fig. 4F**). Reactome pathway enrichment analysis of these miRNA targets identified significant enrichment of pathways related to cellular stress response, various signaling pathways (SCF-KIT, ERBB4, FCERI), and chromatin organization and modification (**Fig. 4F**). Notably, the most significantly enriched pathway was “cellular response to stress” (FDR = 0.000286), suggesting that dysregulation of stress pathways may underlie the association between these miRNAs and EAD. No significant pathways were identified for the two small RNAs (Gly TCC tRF3CCA and miR-1247-5p) consistently downregulated in EAD samples. Together, these findings demonstrate that small RNA profiles differ between EAD and NF groups, regardless of IRI status, suggesting that pre-implantation profiles may be sufficiently informative to aid in predicting EAD.

**Figure 4.**
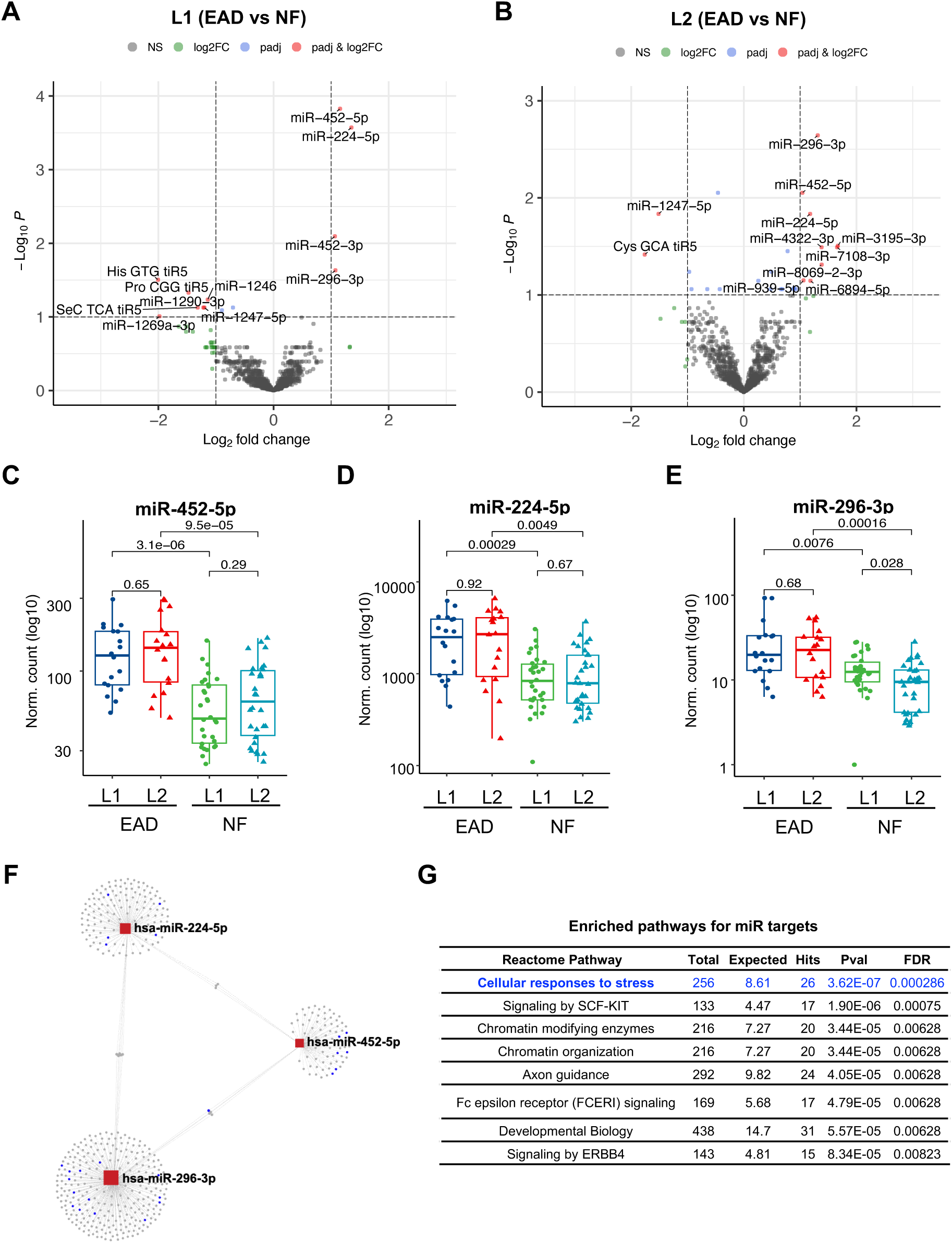
Transplant outcome-associated differences in liver biopsy small RNA expression. **(A)** Volcano plot of differentially expressed small RNAs between early EAD and NF in L1 samples (n = 18 and 30 respectively), identified using DESeq2 (p. adj < 0.1). The x-axis represents log2 fold change (EAD vs. NF), and the y-axis indicates statistical significance (−log10 adjusted p-value). The topmost significantly altered small RNAs are labeled. **(B)** Volcano plot of differentially expressed small RNAs between EAD and NF in L2 biopsy samples (n = 18 and 30 respectively). The topmost significantly altered small RNAs are labeled. **(C – E)** Boxplots of normalized expression (log10 transformed) of representative miRNAs (miR-452-5p, miR-224-5p, miR-296-3p), stratified by transplant outcome (EAD vs. NF) and biopsy timepoint (L1 vs. L2). All three miRNAs are consistently upregulated in EAD compared to NF, independent of time point. Statistical comparisons were performed using Wilcoxon rank-sum tests; p-values are shown above comparisons. **(F)** The lefthand side shows a network visualization of the three differentially expressed miRNAs, where red squares denote the miRNAs and surrounding circular nodes represent their predicted mRNA targets. Target nodes highlighted in blue correspond to genes belonging to the top enriched Reactome pathway, “Cellular responses to stress.” **(G)** The pathway enrichment results generated from the combined miRNA target set (Reactome database, FDR cutoff = 0.01). For each pathway, the total number of annotated genes, expected overlap, observed hits, p-value, and FDR-adjusted significance are shown. “Cellular responses to stress” emerged as the most significantly enriched pathway, consistent with the clustering of blue target nodes in the network plot.

### Pre-transplant liver perfusate is enriched with tRFs and small RNA species

To determine whether liver biopsy-associated tRF and miRNA changes could be captured non-invasively, we analyzed small RNAs in donor liver preservation solution (now referred to as perfusate). Donor livers were submerged in preservation solution and transported on ice prior to transplantation. Immediately before implantation, perfusate was collected and analyzed based on EAD (n = 49) and NF (n = 112) presence (**Fig. S4A**). RNA was extracted from the perfusate and a robust signal corresponding to small RNA size was detected after library construction (**Fig. 5A and Fig. S4B**). The size distribution from the perfusate appears different from the one derived from tissue, highlighting specific enrichment of small RNAs in the perfusate (**Fig. S4B**). As a readily accessible biofluid that does not require tissue biopsy, liver perfusate represents a promising non-invasive source for biomarker discovery. Small RNA sequencing revealed a high abundance of small RNAs in perfusate, with tRFs as the most prevalent class, followed by mitochondrial tRFs (MtRFs) (**Fig. 5B**). Comparisons between perfusates from EAD versus NF donors showed no significant differences in the overall distribution of small RNA classes; tRFs and miRNAs were similarly abundant across both groups (**Fig. 5C).** Pearson correlations revealed moderate to strong relationships between perfusate and both L1 (mean R = 0.70) and L2 (mean R = 0.72) tissue profiles (**Fig. S5A and S5B**), indicating that perfusate partially recapitulates the tissue molecular landscape. Despite the lack of global differences, the consistently high levels of tRFs/miRNAs and other small RNAs in perfusate highlight its potential as a platform for identifying novel RNA biomarkers predictive of donor liver quality and transplant outcomes.

**Figure 5.**
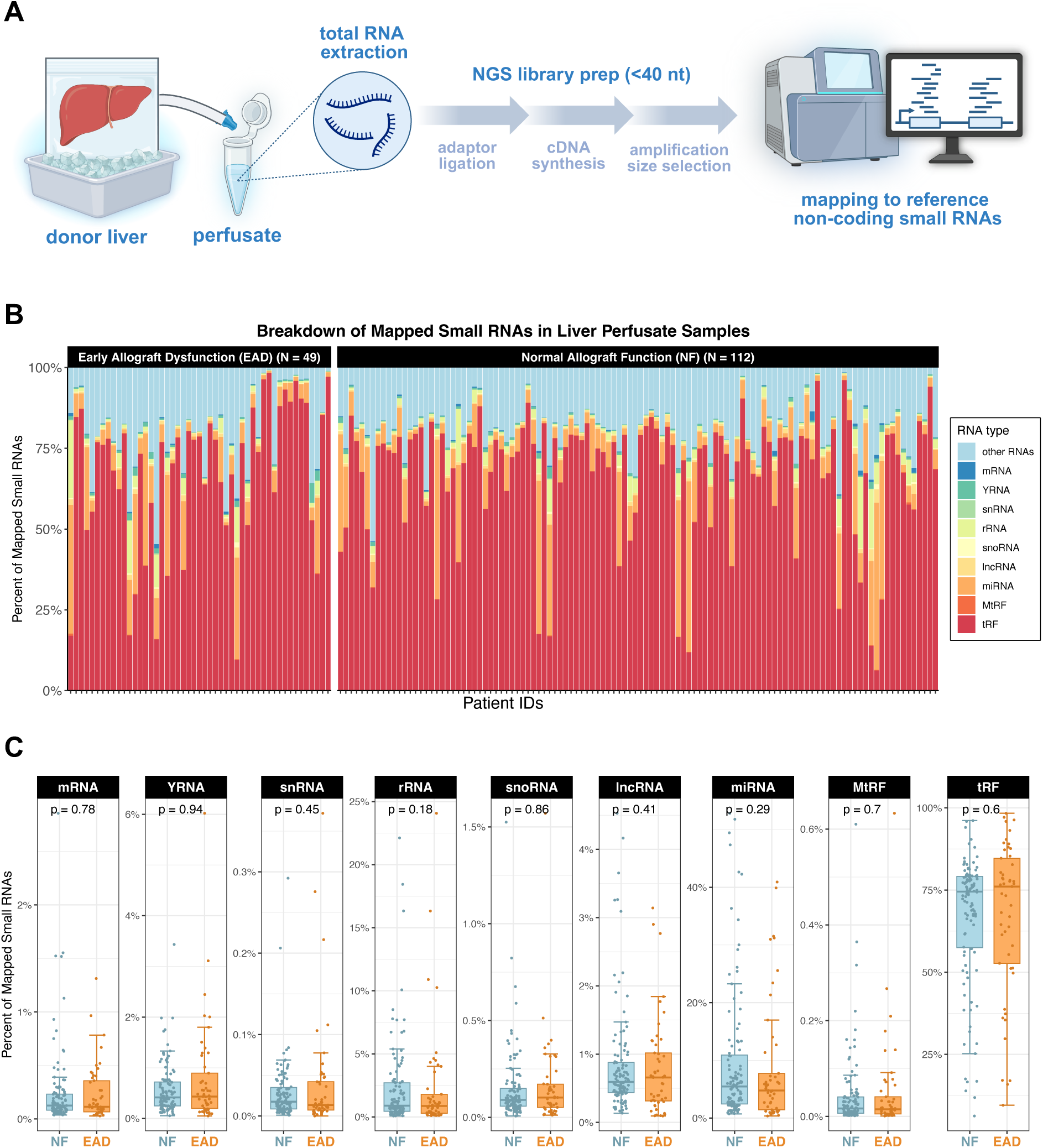
Pre-transplant liver perfusate solution is enriched with small RNAs. **(A)** Donor livers were transported on ice in organ preservation solution prior to transplantation. Immediately before transplantation, the preservation solution (perfusate) was collected, and RNA was isolated for small RNA sequencing**. (B)** Composition of mapped small RNAs in 161 liver perfusate samples, stratified by EAD and NF groups. Across both groups, tRFs represented the predominant RNA species, followed by mitochondrial tRFs (MtRFs). **(C)** Relative abundance of mapped small RNA classes in liver perfusate samples, stratified by NF and EAD groups (n = 49 and 112 respectively). Each boxplot displays the percentage of total mapped small RNAs in each class, with each point representing an individual sample. RNA classes were similarly abundant in both EAD and NF samples, with no statistically significant difference between groups (Wilcoxon test).

### Pre-transplant perfusate small RNAs are analogous to tissue profiles and are differentially expressed in EAD

Given the variable level of tRFs/miRNAs across samples (**Fig. 5C**), we evaluated whether perfusate small RNA composition correlated with donor clinical variables, starting with the donor risk index (DRI), a composite metric that incorporates donor age, cold ischemia time, cause of death, and race (22). We observed that miRNA percentage was significantly positively correlated with DRI (**Fig. 6A**). Similarly, tRF percentage was significancy negatively correlated with DRI (**Fig. 6B**). These correlations suggest that DRI is associated with the overall perfusate small RNA composition. Examination of the individual DRI components indicated that cause of death was the primary contributor to this association (**Fig. S5C**). We next investigated whether pre-transplant perfusate small RNAs could reflect donor liver quality independent from known clinical variables and serve as potential biomarkers for EAD. To do this, we performed differential expression analysis comparing EAD vs. NF groups using DESeq2 with correction for batch and donor risk index (22). Multiple small RNA species showed differential expression in EAD (**Fig. 6C and D, Supplementary Table 2**), suggesting these small RNAs could be potential predictors of EAD independent of donor clinical information. Notably, multiple differentially expressed small RNAs were among the top fifty most highly expressed small RNAs in perfusate samples (**Fig. 6E**). These include Gly GCC/CCC tiR5, Glu CTC/TTC tiR5, Lys CTT tiR5, and miR-26a-5p, all of which were increased in EAD samples, as well as let-7b-5p, which was decreased in EAD samples. Collectively, these findings demonstrate that pre-transplant liver perfusate can serve as a non-invasive source of small RNAs that partially reflect donor liver tissue profiles. The presence and differential expression of perfusate-derived small RNAs further support their potential utility as non-invasive biomarkers for predicting EAD.

**Figure 6.**
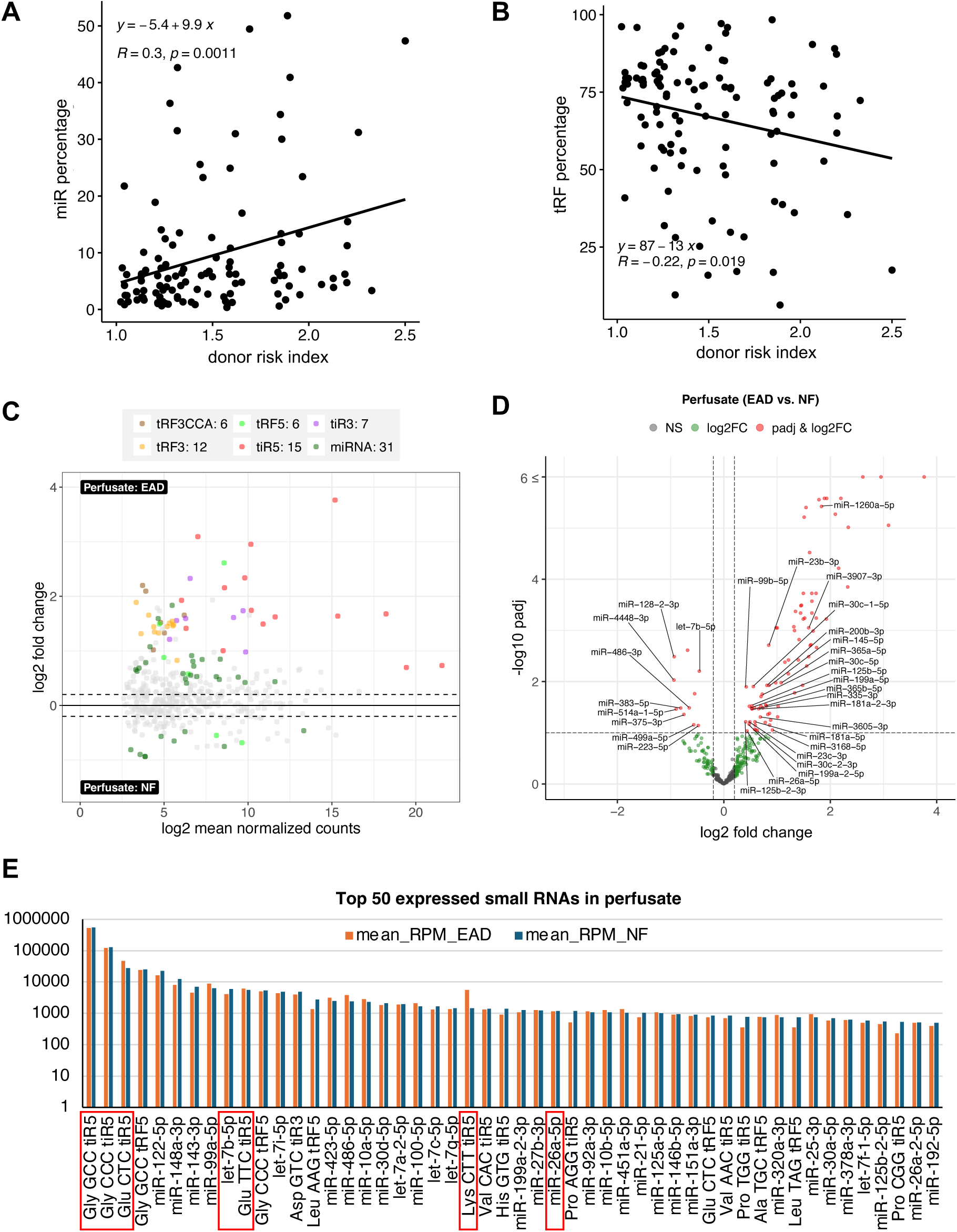
Pre-transplant perfusate small RNAs are differentially expressed based on transplant outcome. **(A)** Scatter plot depicting the relationship between donor risk index (DRI) and the percentage of small RNA sequencing reads mapped to miRNAs in perfusate samples. A significant positive correlation was observed between miRNA percentage and DRI (R = 0.30, p = 0.0011). The DRI incorporates donor age, cold ischemia time, cause of death, and race. **(B)** Scatter plot showing the relationship between DRI and the percentage of small RNA sequencing reads mapped to tRFs. A significant negative correlation was observed between DRI and tRF percentage (R = −0.22, p = 0.019). **(C)** MA plot of the same comparison (EAD vs. NF) in perfusate samples, where the x-axis represents log_2_ mean normalized counts and the y-axis represents log_2_ fold change. Significantly differentially expressed small RNAs (adjusted p-value < 0.1) are color-coded by RNA class (miRNA or tRF subtype). **(D)** Volcano plot showing differential expression of small RNAs in pre-transplant liver perfusate, identified using DESeq2. The x-axis represents log_2_ fold change (EAD vs. NF), and the y-axis shows −log_10_ adjusted p-value. For visualization, all –log10 p-adj values greater than 6 were capped and displayed as 6. Small RNAs meeting significance (adjusted p-value < 0.1) and fold-change thresholds (|log_2_FC| ≥ 0.2) are highlighted, while differentially expressed miRNAs labeled**. (E)** Bar plot showing expression levels (mean RPM) of the top fifty most abundant small RNAs detected in liver perfusate samples, stratified by transplant outcome (EAD, orange; NF, blue). Small RNAs that were significantly differentially expressed in EAD vs. NF comparisons are outlined in red.

### EAD-downregulated perfusate miRNAs improve prediction of early allograft dysfunction

To evaluate whether perfusate small RNAs differentially expressed between EAD and NF could serve as predictive biomarkers, we performed 5-fold cross-validated ROC analyses using multiple feature sets derived from DESeq2 results using 155 patients with available clinical parameters. Predictive performance was compared against using DRI alone. Models incorporating EAD-upregulated perfusate small RNAs did not outperform DRI alone (AUC = 0.665; 95% CI, 0.576–0.754). Notably, upregulated miRNAs performed poorly both alone and in combination with DRI (AUC = 0.596; 95% CI, 0.490–0.701; **Fig. S5D**). In contrast, models based on EAD-downregulated perfusate small RNAs demonstrated improved discriminatory capacity. A combined set of downregulated miRNAs and tRFs achieved an AUC of 0.756 (95% CI, 0.668–0.844) alone and 0.764 (95% CI, 0.677–0.851) when combined with DRI (**Fig. 7A**). Stratification by small RNA class revealed that this improvement was driven primarily by miRNAs. Downregulated perfusate miRNAs alone achieved an AUC of 0.762 (95% CI, 0.675–0.850), which further increased to 0.772 (95% CI, 0.686–0.859) when combined with DRI, representing an 10.7% improvement over DRI alone (**Fig. 7B**). DeLong’s test revealed no significant difference between the miRNA-alone and DRI-alone models (p = 0.106), indicating comparable predictive performance. In contrast, the combined miRNA + DRI model significantly outperformed the DRI-only model (p = 0.033), demonstrating that the addition of these miRNAs resulted in a significant improvement in discriminatory capacity.

**Figure 7.**
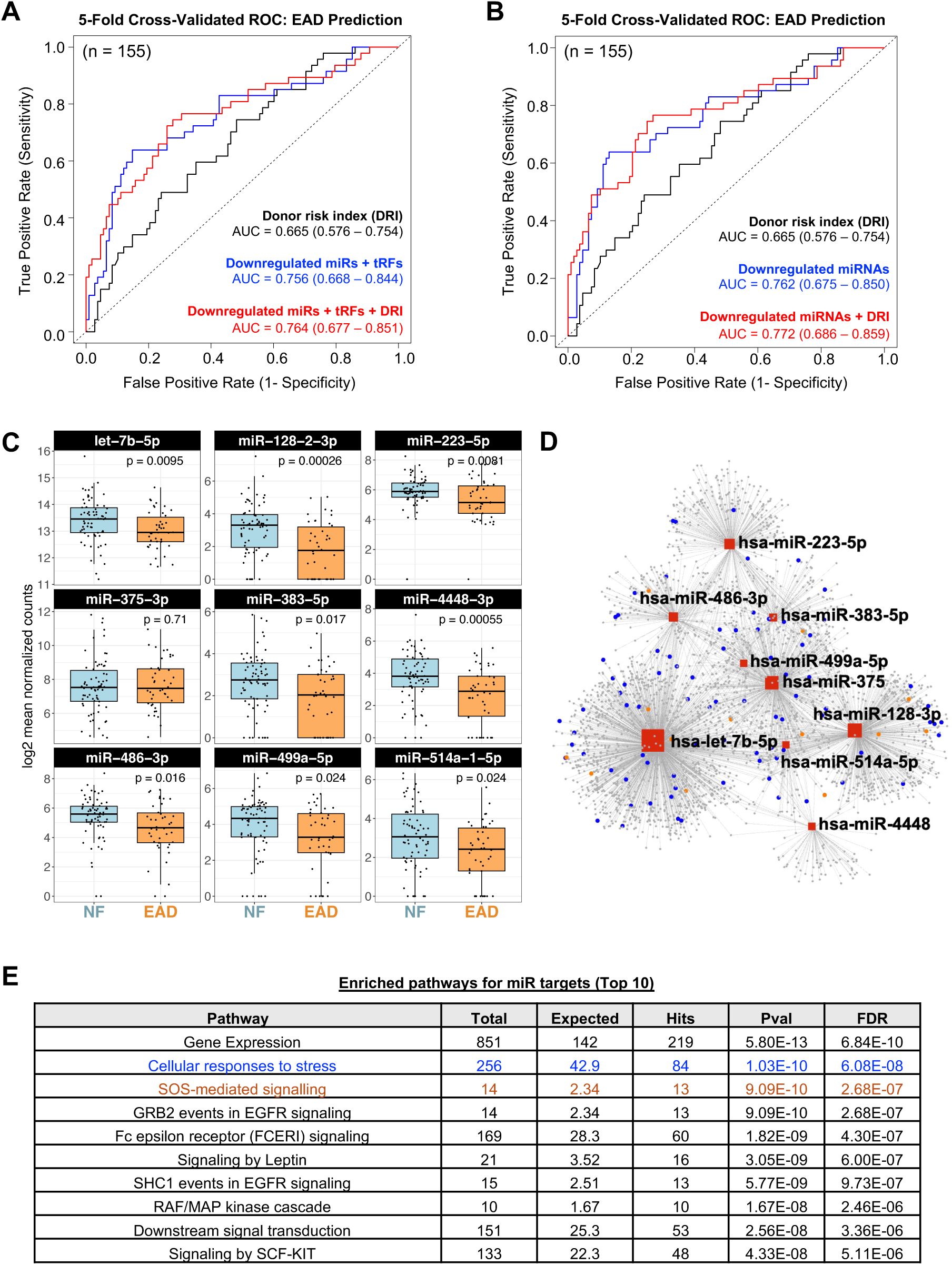
Downregulated perfusate miRNAs candidates improve EAD prediction. **(A)** ROC curves comparing three models for predicting EAD: donor risk index (DRI; black), downregulated perfusate tRFs + miRNAs (blue), and a combined model incorporating both perfusate RNAs and DRI (red). The DRI model includes donor age, cold ischemia time, cause of death, and race. The RNA-only model incorporates expression levels of fourteen perfusate RNAs (miRNAs and tRFs) that were significantly downregulated in EAD compared with NF samples. Model performance, assessed by 5-fold cross-validation, yielded AUCs of 0.665 (DRI), 0.756 (downregulated miRNAs + tRFs), and 0.764 (combined model). Comparisons using DeLong’s test showed no significant differences between the RNA-alone and DRI-alone models (RNA-only vs. DRI, p = 0.13), however the combined model (RNAs + DRI) vs. DRI revealed a significant improvement in AUC (p = 0.047). **(B)** ROC curves comparing DRI alone (black), downregulated perfusate miRNAs (blue), and a combined miRNA + DRI model (red). The miRNA-only model included nine miRNAs significantly downregulated in EAD perfusate samples relative to NF. The miRNA model outperformed DRI alone (AUC 0.762 vs. 0.665), and the combined model further improved discrimination (AUC 0.772), demonstrating that downregulated perfusate miRNAs add predictive value beyond established donor clinical variables. DeLong’s test confirmed the statistically significant difference between the combined model and DRI-only model (p = 0.033; miRNA-only vs. DRI, p = 0.107). **(C)** Boxplots illustrating the expression of the nine EAD-downregulated perfusate miRNAs used in the ROC analysis (panel B). The y-axis shows log₂ normalized counts, with individual points representing each donor perfusate sample. Reported p-values (Wilcoxon test) reflect significant downregulation of these five miRNAs in EAD relative to NF. **(D)** Network visualization of the nine candidate miRNAs (downregulated in EAD vs. NF), where red squares denote the miRNAs and surrounding circular nodes represent their predicted mRNA targets. Target nodes highlighted in blue correspond to genes belonging to the Reactome pathway, “Cellular responses to stress,” while the nodes highlighted in orange correspond to the top enriched pathway “SOS-mediated signaling.” **(E)** Pathway enrichment results generated from the combined miRNA target set (Reactome database, top 10). For each pathway, the total number of annotated genes, expected overlap, observed hits, p-value, and FDR-adjusted significance are shown.

This miRNA model comprised nine miRNAs (let-7b-5p, miR-128-2-3p, miR-223-5p, miR-375-3p, miR-383-5p, miR-4448-3p, miR-486-3p, miR-499-5p, and miR-514a-1-5p) significantly downregulated in EAD perfusate samples (**Fig. 7C**), six of which showed negative correlations with post-transplant day 1 AST and/or ALT levels (**Fig. S6B**). In contrast, downregulated perfusate tRFs did not improve predictive performance beyond DRI (AUC = 0.648; 95% CI, 0.552–0.743) and failed to show significant correlations with early post-transplant liver injury markers (**Fig. S6B and C**). To explore potential biological mechanisms underlying these associations, we performed pathway enrichment analysis of predicted mRNA targets of the nine EAD-downregulated perfusate miRNAs. Network analysis demonstrated substantial overlap in target gene networks among these miRNAs (**Fig. 7C**). Reactome pathway enrichment analysis identified significant enrichment of multiple signaling pathways, including SOS-mediated signaling, FCERI, EGFR, leptin, RAF/MAP kinase, and SCF-KIT signaling pathways (**Fig. 7D and Supplementary Table 3**). Notably, “cellular responses to stress” re-emerged as the most significantly enriched pathway (FDR = 6.08 x 10^−8^), consistent with findings from tissue-based EAD versus NF comparisons (**Fig. 4F** and **Fig. 7D**), while “SOS-mediated signaling” was the second most significantly enriched pathway (FDR = 2.68 × 10⁻^7^). Together, these results demonstrate that EAD-downregulated perfusate miRNAs could serve as predictive biomarkers for early allograft dysfunction independent from established clinical risk metrics. Importantly, these miRNAs can be non-invasively measured in pre-transplant perfusate, highlighting their potential utility for enhancing donor risk stratification prior to transplantation.

## Discussion

In this study, we performed small RNA sequencing on donor liver biopsies from 48 cases and reported abundant tRFs (**Fig. 1**). Analysis of paired post-reperfusion versus pre-implantation biopsies revealed that ischemia-reperfusion injury increased tiR5 and tRF1 species, alongside a decrease in tRF3CCA (**Fig. 2**). Notably, tiR5 was particularly enriched in post-reperfusion L2 tissues from patients who developed EAD, whereas tRF1 upregulation was specific to those with NF graft outcomes (**Fig. 3**). In parallel, comparison of EAD vs NF in L1 and L2 independently highlighted miRNAs and tRFs which are important for EAD pathology (**Fig. 4**). As a proof of concept to use organ preservation solution (perfusate) for non-invasive biomarker discovery, we sequenced perfusate small RNAs from 161 samples and observed abundant tRFs and miRNAs that are likely derived from liver tissues (**Fig. 5**). Comparison of perfusate small RNAs in patients with EAD versus NF identified multiple differentially expressed small RNAs including tRFs and miRNAs (**Fig. 6**). Importantly, a set of nine downregulated perfusate miRNAs can predict post-transplant outcome more effectively than using donor clinical information alone (**Fig. 7**). Our integrated analysis of liver biopsies and perfusate revealed that both compartments contain a shared repertoire of small RNAs, highlighting the potential utility of these small RNAs as non-invasive biomarkers for IRI and post-transplant outcomes.

The observed increase of multiple tiR5 species in L2 (**Fig. 2B-D**) is likely due to the hypoxia and oxidative stress incurred during static cold storage and reperfusion, consistent with the well-established role of tiR5s as stress-induced RNAs, including in hypoxic contexts (4,13). During cellular stress, these tiR5s have been shown to bind to translation initiation factors such as eIF4G/A and displace the eIF4F complex from mRNAs, resulting in global suppression of protein synthesis while permitting selective translation of stress-adaptive transcripts (23). In addition, tiR5s facilitate the assembly of stress granules, sequestering non-essential mRNAs to further conserve resources and prioritize production of proteins critical for survival and recovery (8). Beyond stress adaptation, infection-induced tiR5s have also been implicated in immune activation, particularly through Toll-like receptor 7 interaction (24–26). Given that IRI is characterized as a sterile inflammatory process in which innate immunity plays a central role, the observed tiR5 induction may represent a mechanistic link between cellular stress and immune activation. In particular, the greater tiR5 increase in patients who developed EAD (**Fig. 3B-C**) could indicate more tissue damage or immune activation in those transplants. Conversely, the specific increase in tRF1s upon reperfusion in patients with NF (**Fig. 3B and Fig. S2D**) raises the possibility of a protective role for tRF1 that warrants further mechanistic exploration. Future investigation to identify the molecular targets of these clinically relevant liver tRFs will be needed to reveal their biological functions in the context of IRI stress and organ transplantation (**Fig. 8A**).

**Figure 8.**
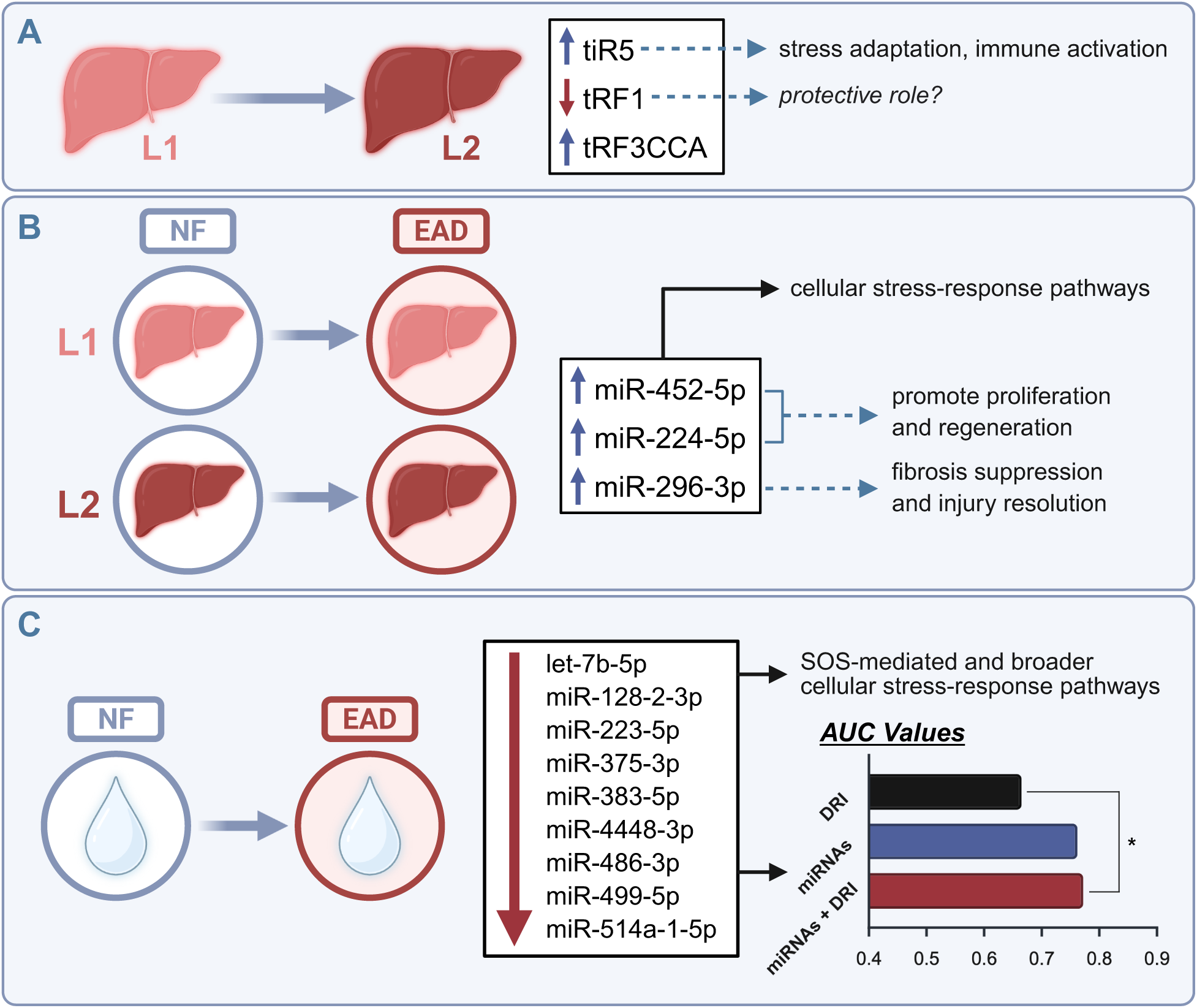
Summary of findings and potential role of tRFs and miRNAs in IRI, EAD and clinical outcome. **A)** Overview of small RNAs differentially expressed in L2 relative to L1 tissue. **B)** Summary of miRNAs and enriched pathways identified in EAD vs NF comparisons across both L1 and L2 tissue. **C)** Summary of nine downregulated perfusate miRNAs with predictive value for EAD and their associated enriched pathways.

Comparisons of EAD versus NF samples in both L1 and L2 tissue cohorts independently identified upregulated miRNAs, including miR-452-5p, miR-224-5p, and miR-296-3p (**Fig. 4A–E**). Target-gene enrichment analysis showed that these miRNAs converge on SOS-mediated and stress-response pathways (**Fig. 4F–G**). Functionally, miR-452-5p has been shown to promote proliferation and cell-cycle progression in hepatocellular carcinoma (27), suggesting a potential role in supporting regenerative responses following injury. Similarly, E2F1-driven induction of miR-224-5p is required for liver cancer progression and invasion (28), indicating that miR-224-5p may enhance cell survival under stress. In contrast, miR-296-3p has been reported to attenuate liver fibrosis (29). Together, these observations suggest a coordinated response in EAD patients in which miR-452-5p and miR-224-5p may promote proliferation and regeneration, while miR-296-3p may contribute to fibrosis suppression and injury resolution (**Fig. 8B**).

The identification of tRFs as dominant small RNAs in perfusate is particularly exciting and demonstrates their potential for use in liquid biopsy monitoring (**Fig. 5**). This is particularly important for evaluating marginal organs and machine perfusion settings, where clinically validated, predictive molecular markers are still lacking. Previously, Verhoeven et al. identified mRNAs and microRNAs in graft preservation solution (perfusate) correlating with early injury (20), while Matton et al. reported that miRNA profiles in normothermic perfusion solution could inform liver viability assessments (21). Both studies were limited to targeted approaches. Darden et al. recently compared miRNA profiles across donor types and recipient outcomes using liver preservation solution and identified distinct miRNA signatures for each donor category, although the study was limited by a small patient cohort (30). Our application of unbiased, next-generation sequencing to perfusate along with tissue small RNA profiling identified clinically relevant miRNAs and tRFs (**Fig. 6 and Fig 7**).

Intriguingly, the set of downregulated miRNAs that enhance EAD prediction (let-7b-5p, miR-128-2-3p, miR-223-5p, miR-375-3p, miR-383-5p, miR-4448-3p, miR-486-3p, miR-499-5p, and miR-514a-1-5p) is enriched in SOS-mediated and broader cellular stress-response pathways (**Fig. 7 and Fig. 8C**). SOS functions as a key upstream activator of Ras proteins, initiating the canonical Ras/Raf/MEK/ERK signaling cascade. This pathway plays a crucial role in promoting the proliferation and survival of hepatocytes and hepatic stellate cells, particularly during cellular stress and liver regeneration (31). Among these miRNAs, let-7b-5p and miR-514a-5p are well-characterized tumor suppressors in hepatocellular carcinoma (32,33). Their reduced expression in EAD may relieve inhibitory constraints on hepatocyte proliferation, thereby supporting early survival responses following graft injury. Furthermore, previous experimental studies have shown that inhibition of miR-128-3p attenuates liver ischemia–reperfusion injury in mice (34), which is consistent with our observation that miR-128-3p is also downregulated in EAD patients. Together, these findings suggest that clinically relevant stress-responsive miRNAs may actively modulate hepatocyte survival pathways during early allograft injury. Targeting or modulating these miRNAs could represent a promising strategy to enhance cellular recovery and improve postoperative outcomes in EAD (**Fig. 8**). In addition, although these small RNAs predict the initial post-transplant outcome, their roles and potential influence in long-term clinical trajectories such as rejection and overall graft lifespan requires further investigation.

Studies of this kind will ultimately enable the transplant community to identify robust small RNA–based biomarkers and accelerate the development of practical detection platforms for real-world application. qPCR-based miRNA and tRF panels, for example, could be implemented during organ procurement or machine perfusion to support rapid, on-site assessment of graft quality. In parallel, emerging CRISPR-based RNA detection technologies, such as the Cas13-driven SHERLOCK system (35), offer highly sensitive, rapid, and field applicable alternatives that could further strengthen intraoperative decision making. Moreover, recent advances in serum and plasma small RNA detection for cancer associated miRNAs (36,37) provide a strong technical foundation that is readily translatable to organ transplantation workflows.

While these opportunities are compelling, several technical considerations inherent to small RNA sequencing must be acknowledged. Small RNA-seq library preparation is prone to sequence-specific biases, including differences in adapter ligation efficiency, GC content, and secondary structure, which can lead to over– or under-representation of certain small RNA species. These biases, however, are well described and can be minimized through standardized protocols and appropriate normalization strategies, both of which were applied in our analysis. Although variation in sequencing depth can affect detection of very low-abundance miRNAs, the depth achieved here was sufficient to reliably characterize the predominant small RNA populations in all sample types. Due to limited RNA availability, particularly in perfusate samples, we were unable to perform additional qPCR validation, and therefore our findings rely on sequencing-derived quantification alone. As such, while the identified small RNA signatures are biologically plausible and reinforced by pathway-level analyses, they should be validated in future studies using independent cohorts and targeted assays.

## Conclusion

Despite the technical limitations, this study provides the first transcriptome-wide characterization of small RNAs in both human liver tissue and donor organ perfusate during transplantation. We demonstrate that specific tRFs and miRNAs are closely linked to reperfusion and EAD. The prevalence of tRFs and miRNAs in perfusate highlights their potential as non-invasive biomarkers, bridging molecular insights from tissue with clinically accessible diagnostics. By uncovering both shared and compartment-specific signatures, our findings lay a foundation for future validation in larger cohorts and under machine perfusion, advancing systems-level small RNAs toward clinical application for improved pre-operative organ assessment and peri-operative transplant monitoring.

## Methods

### Patient recruitment and specimen collection

Liver tissue and perfusate samples were obtained from adult orthotopic liver transplant (OLT) recipients (≥18 years old) at Methodist University Hospital after obtaining informed consent. All procedures were conducted under protocols approved by the University of Tennessee Health Science Center Institutional Review Board (IRB#20-07742-XM and IRB#20-07640-XP). Patients were followed for 2 years after transplant. EAD and NF cohorts were defined based on Day 1 AST values based on Olthoff et al EAD definition. Needle biopsy specimens collected pre-implantation (L1) and post-reperfusion (L2) from the same patients were immediately immersed in RNA*later* (Thermo Fisher Scientific, Cat. #AM7021) and incubated at 4 °C overnight to ensure complete permeation. RNA*later* was then removed, and the samples were stored at –80 °C until RNA extraction.

Perfusate samples were collected at the end of organ procurement concurrently with pre-implantation liver (L1) biopsies. The solutions were centrifuged at 2,400 rpm for 30 minutes at room temperature to remove debris. The resulting supernatant was aliquoted and stored at –80 °C until RNA isolation.

Donor livers were transported on ice in the perfusate solution. Average cold ischemia time is 4.5 hours. The perfusate samples were collected in the operation room at the back table before organ was prepped for the transplant and directly placed on ice and transferred to fridge. The perfusate samples were centrifuged and debris was removed. Lastly, the samples were kept at –80°C until RNA isolation. If the perfusate sample had to stay in the fridge overnight and were not centrifuged and frozen on the same they, they were not used in this study.

### RNA isolation and small RNA library preparation

Total RNA was extracted from needle biopsies using the Direct-zol RNA MicroPrep Kit (Zymo Research, Cat. #R2062) according to the manufacturer’s instructions, including on-column DNase I treatment to remove genomic DNA contamination. For perfusate samples, RNA was isolated from 500 μl of solution using the same kit, following the manufacturer’s protocol for liquid samples, and eluted in 15 μl of RNase-free water. RNA concentrations were ranging 50-300 ng/ μl. Purified RNA was stored at –80 °C, and repeated freeze–thaw cycles were avoided to minimize RNA degradation. RNA quality was evaluated using QIAxcel, and each tissue sample exhibited an RNA integrity score (RIS) greater than 6, consistent with high-quality RNA. Perfusate RNA is composed primarily of small RNAs and lacks sufficient intact ribosomal RNA, making it unsuitable for quality assessment using QIAxcel or the Illumina Bioanalyzer. Perfusate RNA concentrations were extremely low and impossible to detect using Qubit RNA HS Assay. Small RNA libraries were prepared using the NEBNext Multiplex Small RNA Library Prep Set for Illumina (New England Biolabs, Cat. #E7330) following the manufacturer’s instructions. For each sample, 100 ng of tissue RNA or 5 μl of perfusate RNA was used as input. Indexing was performed using the NEBNext Multiplex Oligos for Illumina, with PCR amplification carried out for 15 cycles for tissue RNA libraries and 18 cycles for perfusate RNA libraries. Amplified products were resolved on an 8% polyacrylamide gel (Thermo Fisher Scientific, Cat. #EC6215BOX), and target bands were excised and purified. Equal amounts of libraries were pooled and sequenced on an Illumina NextSeq 500 platform, generating approximately 10 million 75 bp single-end reads per sample.

The samples were divided into carefully planned batches. Tissue RNA libraries were prepared in three batches and sequenced in three corresponding batches. For each patient, L1 and L2 samples were always processed within the same batch, and every batch included both EAD and NF samples. Perfusate samples were similarly organized so that each batch contained a mix of EAD and NF samples, minimizing potential batch effects.

### Small RNA-seq data analysis

Raw small RNA sequencing reads were trimmed to remove adapter sequences using Cutadapt (v5.1). Adapter-trimmed reads were then mapped and annotated using Unitas (v1.7.3), a specialized tool for comprehensive small RNA annotation similar to previously described (38). Briefly, unitas regular mode allows 1 internal mismatch and maps reads first to human miRbase Release 22 (39), then to other small RNA sequences including genomic tRNA database (40), SILVA rRNA database Release 132 (41) and Ensembl Release 97. In the case of multi-mapping, a read was counted as fraction distributed equally to avoid duplicate counts. For liver biopsy samples, read count matrices generated by Unitas were imported into DESeq2 (v1.44.0) in R for normalization and differential expression analysis. Prior to analysis, RNAs with zero counts in more than 80% of samples were excluded to reduce low-abundance noise. Data visualization was performed in R using ggplot2 (v3.5.1) and ggpubr (v0.6.0). For perfusate samples, read count matrices generated by Unitas were imported into R and normalized using DESEq2 with batch correction and DRI in the model. For L2 vs L1 comparisons, DESeq2 design includes patient for paired analysis, all DESeq2 output is included in **Supplementary Table 1**. For EAD vs NF comparisons, DESeq2 design includes batch effect and DRI scores, all DESeq2 output is included in **Supplementary Table 2**. The corresponding sample number and significance threshold is indicated in the Figure legends. Data visualization was carried out in R with ggplot2 (v3.5.1) and ggpubr (v0.6.0).

### Clinical correlation and ROC analysis

Donor clinical variables were correlated with perfusate small RNA abundance using Pearson correlation in R. Correlation matrices were visualized with the corrplot package (v0.92). Donor Risk Index (DRI) scores were calculated according to the equation defined by Feng et al. (22), which incorporates seven donor or graft characteristics that independently predict graft failure: donor age, donor race, donor height, cause of death, donation after cardiac death, split/partial graft status, and cold ischemia time. ROC curves were generated in R using the ROCR package (v1.0-11). Models were trained using 5-fold cross-validation, in which the dataset was randomly partitioned into five equal subsets; four folds were used to train a logistic model, and the remaining fold served as a held-out test set. This process was repeated five times so that each fold was used once for testing. Reported AUC values represent the mean across give folds. Comparisons between AUCs were performed using DeLong’s test in the pROC package (v1.18.5), to assess whether performance differences between models were statistically significant.

### miRNA target prediction and pathway analysis

microRNA target network analysis was performed using miRNet2.0 (42). microRNA targets are predicted by miRTarBase v9.0 (43) and pathway enrichment calculated using hypergeometric test against Reactome pathways (44).

## Declarations

### Ethics approval and consent to participate

This study was conducted in accordance with the ethical standards of the University of Tennessee Health Science Center Institutional Review Board, which approved all protocols (IRB#20-07742-XM and IRB#20-07640-XP). Liver tissue and perfusate samples were obtained from adult orthotopic liver transplant (OLT) recipients (≥18 years old) at Methodist University Hospital after written informed consent was obtained from all participants.

### Availability of data and materials

The small RNA-seq data has been deposited to GEO (GSE311219 and GSE311223).

### Authors’ contributions

C.W., G.R., and M.K. helped with patient sample processing, RNA isolation and small RNA library preparation. M.D. and Cem K. coordinated sample collection, storage and contributed to study design. J.E., J.V., and C.E. coordinated patient recruitment and sample collection at the Methodist Hospital. M.A., K.M. and Z.S. performed data analysis on RNA-seq. Canan K. and Z.S. supervised the study and wrote the manuscript with the help of M.A and K.M. All authors read and approved the final manuscript.

## Acknowledgements

We would like to thank the Methodist University Hospital transplant team for their assistance with patient recruitment, sample and clinical data collection. We are deeply grateful to the patients and their families for their participation and generous donations to this research. We also would like to acknowledge our colleagues at the Transplant Research Institute for their helpful discussions and ongoing support. We would like to thank IT Research Computing Team at the University of Alabama at Birmingham for NGS support.

