## Supplementary Figures for "Comparative Landscape of Small RNAs in Tissue and Liquid Biopsies for Liver Transplant Outcomes"

### Supplementary Figure 1

A

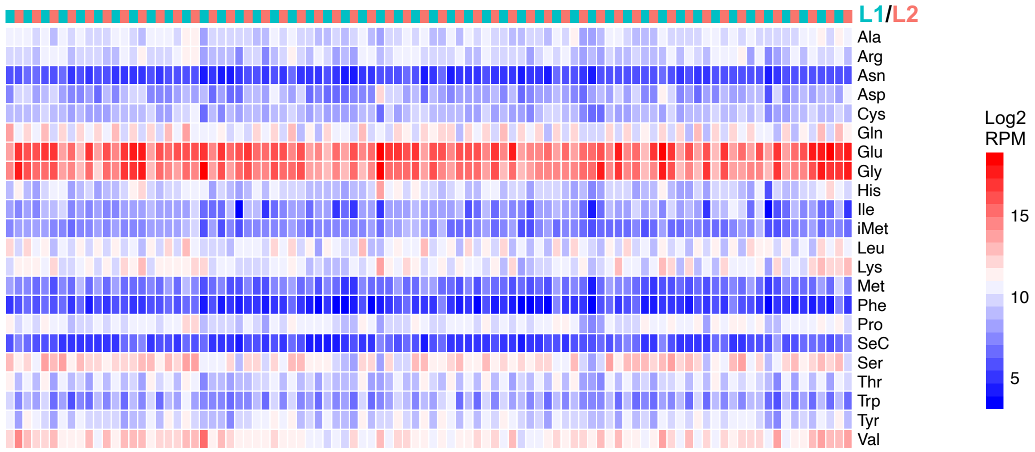


B

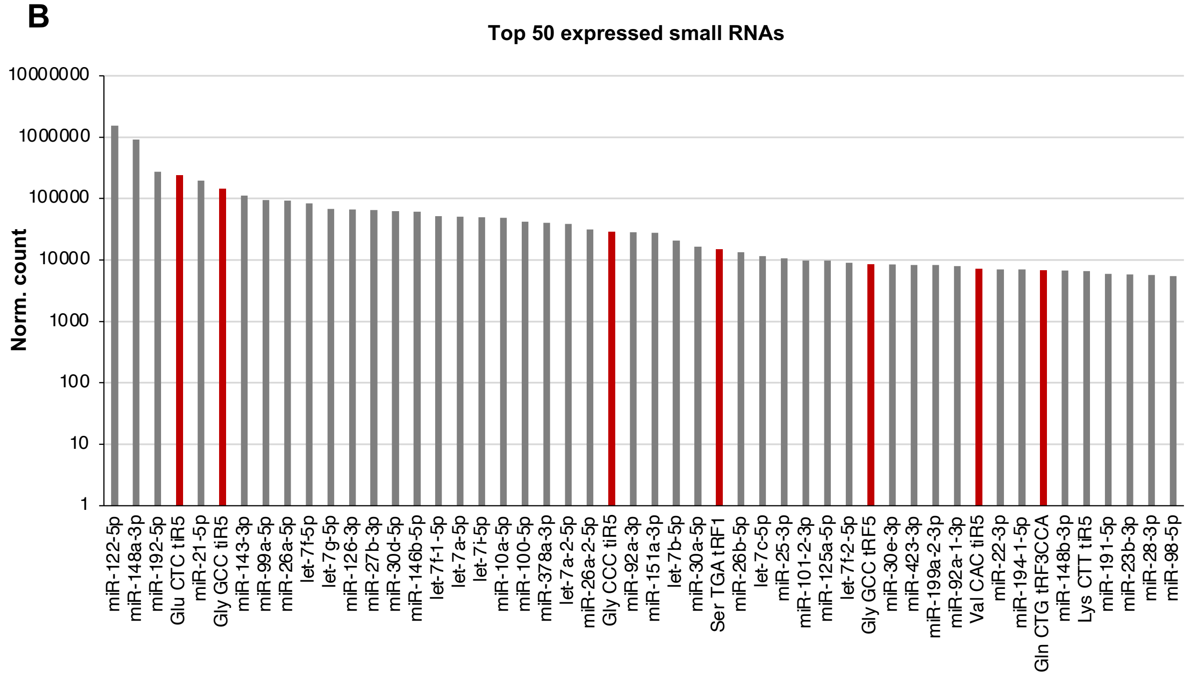

**Supplementary Figure 1. Abundance of tRFs in donor liver transplant biopsy samples.**

**(A)** Heatmap showing tRF expression across liver biopsy samples (L1 and L2), organized by the amino acid carried by the parental tRNA. Expression values are displayed as log2 RPM. tRFs derived from Glu- and Gly-carrying tRNAs are highly enriched relative to other amino acid isotypes**. B)** Bar plot displaying the expression (DESeq2 normalized counts) of the top fifty most highly expressed small RNAs in donor liver biopsy samples. These top fifty genes consist entirely of miRNAs (gray bars) and tRFs (red bars).

### Supplementary Figure 2

**A**

| Group | EAD | NF | p value |
|---|---|---|---|
| **Mean INR (Day 1 post transplant)** | 2.10 | 2.03 | N.S. |
| **Mean AST (Day 7 post transplant)** | 130.3 | 70.7 | N.S. |
| **Mean ALT (Day 7 post transplant)** | 235.4 | 121.9 | 0.017 * |
| **Mean INR (Day 7 post transplant)** | 1.22 | 1.16 | N.S. |

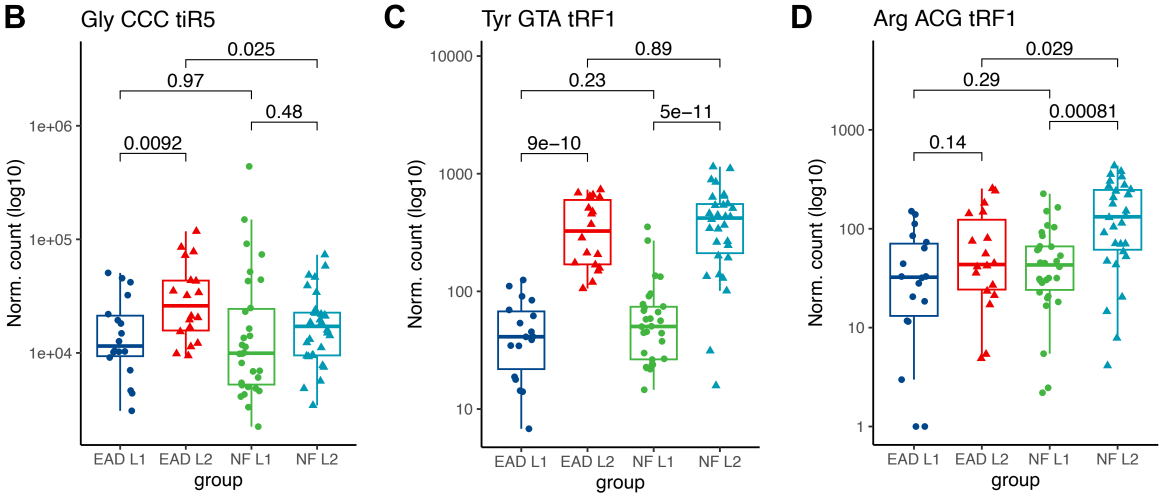


**E**

**DOWN-regulated in L2 (post) vs L1 (pre)**

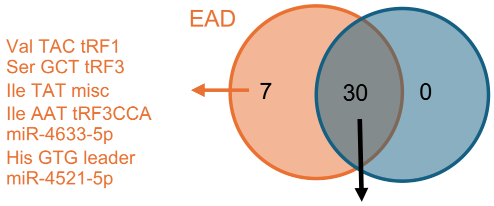


Gln CTG tRF3CCA, Gln CTG tRF3, Gln TTG tRF3CCA, Gln TTG tRF3, Ser CGA tRF3, Ser CGA tRF3CCA, miR-5100-3p, Leu CAA tRF3CCA, Pro AGG tRF3, Pro TGG tRF3, Leu CAA tRF3, Pro CGG tRF3, Thr TGT tRF3CCA, Ser AGA RF3CCA, Trp CCA tRF3CCA, Ser TGA tRF3CCA, Pro AGG tRF3CCA, Pro TGG tRF3CCA, Pro CGG tRF3CCA, Ser AGA tRF3, Tyr GTA tRF3CCA, Leu CAG tRF3CCA, Thr TGT tRF3, Trp CCA tRF3, Ser TGA tRF3, Ser GCT tRF3CCA, Thr CGT tRF3CCA, Leu AAG tRF5, Arg TCG tRF3CCA, Leu TAG tRF5

**F**

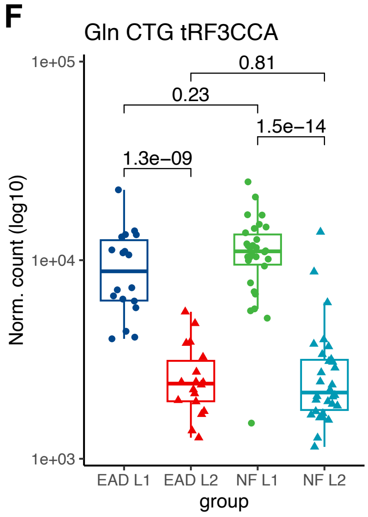

**Supplementary Figure 2. Additional outcome-specific IRI-associated tRF alterations.**

**(A)** Table of additional post-transplant clinical parameters, stratified by patient transplant outcome (EAD or NF). **(B—D, F)** Boxplots showing the normalized expression (log10 transformed) of selected tRF species across outcome groups (EAD or NF) and biopsy timepoints (L1 or L2). The selected tRF species represent examples of changes between L2 and L1 samples that are specific to EAD, NF, or shared by both groups. **(B)** Gly-CCC-tiR5 is significantly upregulated in L2 compared to L1 biopsies, exclusively within EAD samples. **(C)** Try-GTA-tRF1 is significantly upregulated in L2 across both EAD and NF samples**. (D)** Arg-ACG-tRF1 is significantly upregulated in L2 samples only within the NF group. **(E)** Differential expression analysis of L2 vs. L1 samples was performed separately within EAD and NF groups. The Venn diagram displays small RNAs downregulated in L2 relative to L1 for each group. In the EAD group, uniquely downregulated RNAs included multiple tRF subtypes and miRNA species. NF samples exhibited no uniquely downregulated RNAs, with all L2-downregulated species overlapping with those in the EAD group, the majority being tRF3CCA species. **(F)** Gln-CTG-tRF3CCA is downregulated in the L2 sample across both EAD and NF groups, exemplifying the overlapping downregulated tRF3CCA species presented in panel E.

Supplementary Figure 3

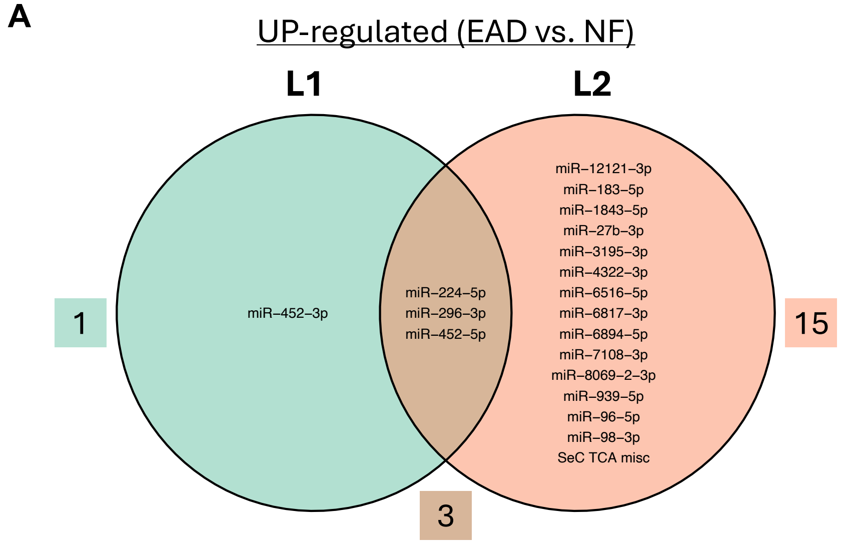


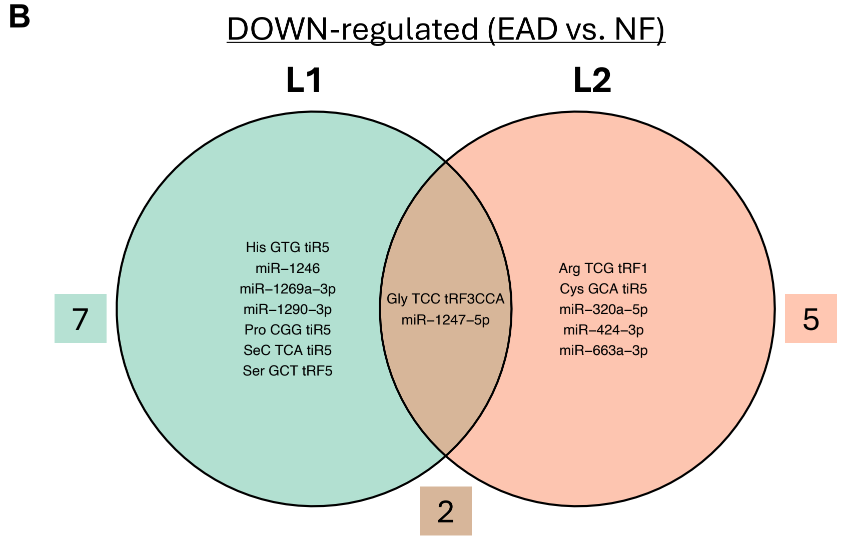

**Supplementary Figure 3. Differential expression of small RNAs between EAD and NF groups.**

**(A)** and **(B)** Differential expression analysis (DESeq2) was performed separately within the L1 and L2 sample sets, comparing EAD vs. NF transplant outcomes. **(A)** Venn diagram showing small RNAs upregulated in EAD compared to NF at either the L1 or L2 timepoint. Three miRNA species were commonly upregulated across both timepoints. **(B)** Venn diagram showing small RNAs downregulated in EAD compared to NF at either the L1 or L2 timepoint. One tRF3CCA fragment and one miRNA were commonly downregulated across both timepoints.

### Supplementary Figure 4

**A**

**Perfusate sample summary (n = 161)**

| Group | EAD | NF | p value |
|---|---|---|---|
| Number of patients | 49 | 112 | |
| Recipient mean age | 58.7 | 58 | N.S. |
| Recipient gender (F/M) | 15/34 | 42/68 | N.S. |
| Donor mean age | 45.6 | 39.8 | 0.011 * |
| Donor gender (F/M) | 14/35 | 55/57 | 0.024 * |
| Mean CIT (hours) | 4.56 | 4.51 | N.S. |
| Mean AST (Day 1 post transplant) | 4644 | 826 | 2.683e-10 *** |
| Mean ALT (Day 1 post transplant) | 1746 | 458 | 7.382e-11 *** |
| Mean INR (Day 1 post transplant) | 1.87 | 1.83 | N.S. |
| Mean AST (Day 7 post transplant) | 86 | 86.5 | N.S. |
| Mean ALT (Day 7 post transplant) | 211 | 123 | 8.385e-06 *** |
| Mean INR (Day 7 post transplant) | 1.20 | 1.18 | N.S. |

**B**

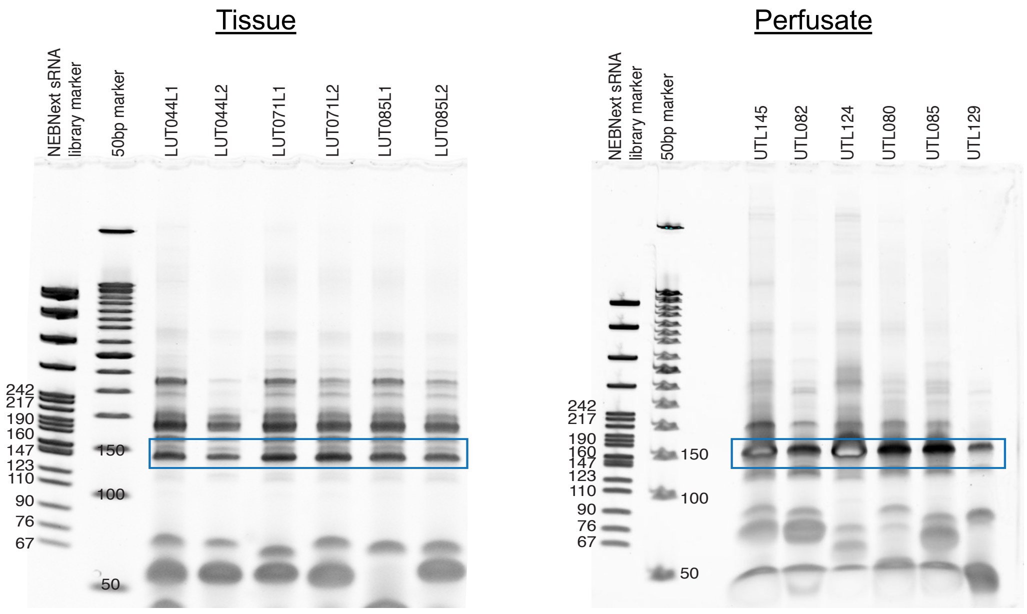

**Supplementary Figure 4. Clinical cohort and cDNA library characteristics.**

**(A)** Summary of demographic and clinical characteristics for the cohort from which perfusate samples were obtained (n = 161), stratified by outcome group (EAD or NF). **(B)** Representative PAGE gels showing small RNA cDNA library preparations from tissue biopsies (left) and perfusate samples (right). Fragments of ~150 bp (highlighted by blue box) were excised and used for small RNA sequencing.

### Supplementary Figure 5

**A**

Pearson's correlation between perfusate and tissue small RNAs

| **Patient:** | LUT005 | LUT007 | LUT023 | LUT024 | LUT039 | LUT010 | LUT045 | LUT072 | LUT046 | LUT004 | LUT014 | LUT015 | LUT044 | LUT085 | LUT088 | LUT099 | LUT052 | LUT057 | 0 |
|---|---|---|---|---|---|---|---|---|---|---|---|---|---|---|---|---|---|---|---|
| **Perfusate ~ L1** | 0.69 | 0.70 | 0.63 | 0.73 | 0.69 | 0.71 | 0.68 | 0.75 | 0.73 | 0.74 | 0.67 | 0.76 | 0.68 | 0.71 | 0.75 | 0.75 | 0.74 | 0.69 | 0.5 |
| **Perfusate ~ L2** | 0.73 | 0.70 | 0.71 | 0.74 | 0.76 | 0.74 | 0.75 | 0.75 | 0.79 | 0.76 | 0.68 | 0.64 | 0.70 | 0.73 | 0.76 | 0.77 | 0.77 | 0.75 | 1 |

| **Patient:** | LUT061 | LUT070 | LUT073 | LUT048 | LUT051 | LUT083 | LUT087 | LUT106 | LUT065 | LUT077 | LUT084 | LUT105 | LUT089 | LUT111 | LUT117 | LUT127 | LUT038 | LUT098 | 0 |
|---|---|---|---|---|---|---|---|---|---|---|---|---|---|---|---|---|---|---|---|
| **Perfusate ~ L1** | 0.85 | 0.72 | 0.70 | 0.66 | 0.67 | 0.63 | 0.59 | 0.68 | 0.75 | 0.65 | 0.76 | 0.62 | 0.56 | 0.69 | 0.74 | 0.76 | 0.73 | 0.68 | 0.5 |
| **Perfusate ~ L2** | 0.82 | 0.66 | 0.68 | 0.72 | 0.70 | 0.67 | 0.65 | 0.67 | 0.79 | 0.69 | 0.73 | 0.69 | 0.67 | 0.71 | 0.76 | 0.77 | 0.74 | 0.73 | 1 |

**B**

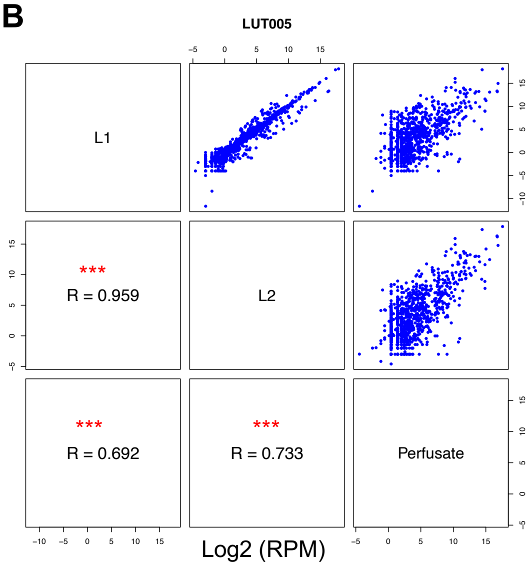


**C**

Correlation between clinical variable and small RNA composition

| | **miR percentage** | **tRF percentage** |
|---|---|---|
| Donor risk index | R = 0.3, p = 0.0011 | R = -0.22, p = 0.019 |
| Donor age | R = 0.17, p = 0.062 | R = -0.049, p = 0.6 |
| CIT | R = 0.13, p = 0.18 | R = -0.15, p = 0.11 |
| Cause of death | R = 0.28, p = 0.0019 | R = -0.29, p = 0.0014 |
| Donor race | R = 0.071, p = 0.45 | R = -0.06, p = 0.53 |
| Donor type | R = 0.11, p = 0.25 | R = -0.097, p = 0.3 |

**D**

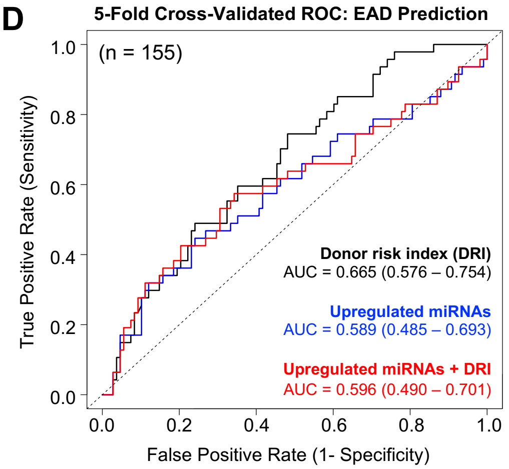

**Supplementary Figure 5. Donor liver perfusate and tissue small RNA profiles are correlated.**

**(A)** Pearson correlation coefficients between perfusate and paired donor liver biopsy samples (L1 and L2). **(B)** Representative pairwise scatterplots of small RNA expression (log2 RPM) in patient LUT005 demonstrate a strong correlation between L1 and L2 biopsy samples (R = 0.959), and moderate correlations between perfusate and L1 (R = 0.692) or L2 (R = 0.733) biopsies**. (C)** Table summarizing correlations between donor clinical variables and perfusate small RNA composition. Pearson correlation coefficients (R) and associated p-values are shown for the percentage of reads mapping to miRNAs (middle column) and tRFs (right column). **(D)** ROC curves comparing three models for predicting EAD: donor risk index (DRI; black), upregulated perfusate miRNAs (blue), and a combined model incorporating both perfusate miRNAs and DRI (red). The DRI model includes donor age, cold ischemia time, cause of death, and race. The miRNA-only model incorporates expression levels of all perfusate miRNAs that were significantly upregulated in EAD compared with NF samples. Model performance, assessed by 5-fold cross-validation, yielded AUCs of 0.665 (DRI), 0.589 (upregulated miRNAs), and 0.596 (combined model). Comparisons using DeLong's test showed no significant differences relative to the DRI model (miRNA-only vs. DRI, p = 0.235; combined vs. DRI, p = 0.231).

Supplementary Figure 6

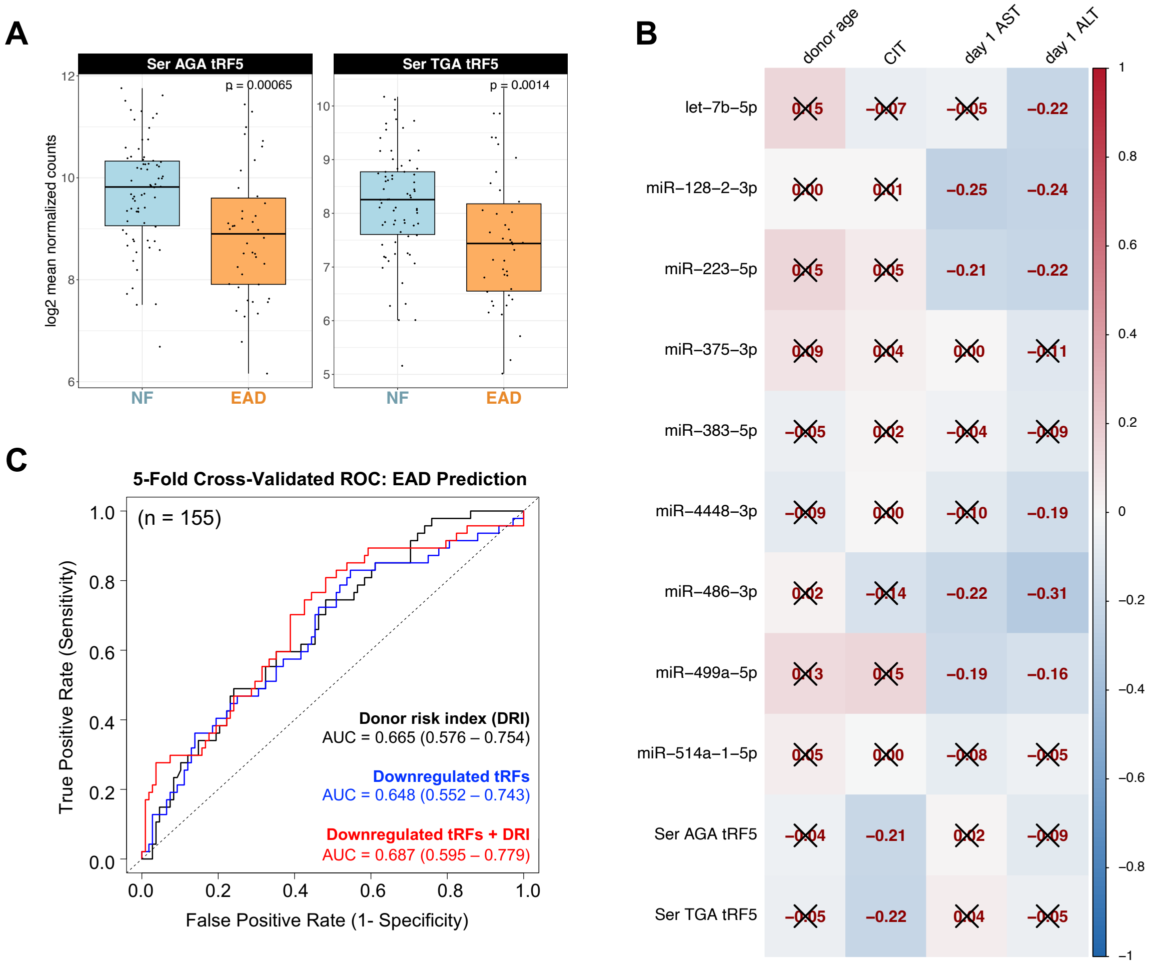

**Supplementary Figure 6. Extended analysis of downregulated perfusate RNA candidates.**

**(A)** Boxplots of all downregulated perfusate tRFs (DESeq2 EAD vs. NF). The y-axis shows $\log_2$ normalized counts, with individual points representing each donor perfusate sample. Reported p-values were determined using Wilcox tests. **(B)** Heatmap showing correlation coefficients (Pearson's R) between downregulated perfusate miRNAs/tRFs and clinical variables. "X" symbols indicate non-statistically significant correlation coefficients (p-value $\geq 0.05$). Six of the nine miRNA candidates showed significant negative correlations with day 1 post-transplant AST and/or ALT values. **(C)** ROC curves comparing DRI alone (black), downregulated perfusate tRFs (blue), and a combined tRF + DRI model (red). The tRF-only model includes two tRFs that were significantly downregulated in EAD perfusate samples relative to NF. Model performance, assessed by 5-fold cross-validation, yielded AUCs of 0.665 (DRI), 0.648 (downregulated tRFs), and 0.687 (combined model); comparisons using DeLong's test showed no significant differences relative to the DRI model (tRF-only vs. DRI, p = 0.783; combined vs. DRI, p = 0.464).
